# Hybrid Transport-of-Intensity and Polarization Differential Phase Contrast for Extended Spatial-Frequency Phase Imaging

**DOI:** 10.64898/2026.09.28.754925

**Authors:** Fraser Montandon, Jasmin Knopp, Caron A. Jacobs, Fred Nicolls

## Abstract

The Transport-of-Intensity Equation (TIE) and polarization differential phase contrast (pDPC) exhibit transfer characteristics that limit their use as standalone quantitative phase imaging modalities. TIE provides stable recovery of low spatial frequencies but attenuates fine structure, while pDPC enhances phase gradients but is intrinsically insensitive near zero frequency. We present a hybrid reconstruction framework that combines a single-shot pDPC measurement with an in-focus–defocused TIE image pair and fuses the resulting phase estimates in the Fourier domain using frequency-selective weighting. The illumination geometry is co-designed to enforce angular separation between modalities, enabling high-frequency phase-gradient acquisition in a single exposure while preserving low-frequency sensitivity through low-numerical-aperture symmetric illumination. The proposed method requires three intensity measurements without requiring interferometric stability or sequential asymmetric illumination. Experimental results demonstrate improved phase fidelity across spatial frequencies, reduced halo artifacts, and enhanced recovery of both smooth and fine structural features. This approach provides a compact and robust method that moves towards extended spatial-frequency phase imaging for dynamic and resource-constrained applications.

## 1. Introduction

Quantitative phase imaging (QPI) provides label-free measurement of optical path length (OPL) variations in transparent specimens, enabling intrinsic contrast in samples that are weakly absorbing or scattering under brightfield illumination [1]. By reconstructing the complex transmitted optical field, QPI yields quantitative phase maps linked to an object’s refractive index (RI) and thickness [2]. These maps support downstream analyses including drug-organism interactions [3], growth monitoring [4, 5], virtual staining [6], cellular diversity and classification [7, 8], viability assessment [9], morphological characterization [1, 10, 11], and segmentation [1]. Owing to its non-invasive nature and independence from exogenous labels or high-intensity fluorescence excitation, QPI enables live-cell and low-perturbation imaging.

QPI encompasses methods that recover phase from intensity measurements and they can be broadly classified by illumination coherence. Digital holography (DH) employs coherent illumination and interferometric detection [12]. Inline DH offers a compact optical configuration but suffers from twin-image artifacts, whereas off-axis DH mitigates these via spatial-frequency separation at the expense of increased optical complexity, reduced spatial bandwidth, and stricter alignment. In both cases, coherent laser illumination introduces speckle and other coherence-induced artifacts that may degrade phase fidelity and limit performance for live or dynamic samples. Partially coherent methods, including differential phase contrast (DPC) [13, 14] and Fourier ptychography (FP) [15], avoid many coherence-related artifacts and can achieve an effective bandwidth approaching twice that of coherent systems with the same objective numerical aperture (NA). However, these methods remain intrinsically band-limited. DPC reconstructs phase gradients via asymmetric illumination but exhibits poor sensitivity to low spatial frequencies. Accurate phase recovery typically relies on the weak object approximation (WOA) [16] and careful matching between illumination and objective NAs.

Hybrid phase imaging strategies aim to overcome these limitations by combining reconstruction modalities. Integrating off-axis DH with FP extends spatial-frequency coverage and improves phase recovery, but retains interferometric acquisition [17], with increased optical complexity, coherence requirements, and sensitivity to mechanical instability. Non-interferometric hybrids that combine FP with transport-of-intensity equation (TIE) constraints have also been proposed [18]; however, they typically require multi-shot FP acquisitions, leading to higher photon dose, longer acquisition times, and increased computational cost. Recently, combining DPC with the TIE has demonstrated improved phase recovery under partially coherent illumination [19]. TIE provides reliable recovery of low-spatial-frequency phase under regularization and reduces nonlinear phase effects arising from WOA violations, while DPC provides high-spatial-frequency phase-gradient contrast. However, existing DPC–TIE implementations typically require multiple sequential asymmetric illuminations in addition to defocused measurements. This increases susceptibility to motion artifacts and limits applicability to dynamic specimens. These limitations motivate an extended spatial frequency QPI framework that minimizes acquisition count while avoiding interferometry and stringent illumination–objective NA matching constraints.

In this work, we introduce hybrid polarization differential phase contrast (hpDPC), a QPI framework that combines polarization differential phase contrast (pDPC) [20] with TIE-based phase recovery. Unlike prior hybrid approaches that combine independently acquired modalities, hpDPC co-designs the illumination geometry and Fourier-domain fusion to promote complementary spatial-frequency responses between pDPC and TIE. The framework integrates polarization-encoded asymmetric illumination for pDPC with a coaxial, NA-limited illumination channel for TIE within a unified hpDPC illumination module. pDPC uses polarization multiplexing and a polarization-sensitive camera to encode multiple asymmetric illumination states in a single exposure, enabling snapshot recovery of high-spatial-frequency phase gradients. Lower frequency components are recovered via TIE from a focused–defocused image pair acquired under symmetric illumination. By structurally separating high-and low-frequency phase transfer between modalities, hpDPC relaxes illumination–objective NA matching and reduces sensitivity to nonlinear phase effects beyond the WOA. Our framework requires only three measurements (one multiplexed pDPC acquisition and two TIE), eliminating sequential asymmetric illumination, and is implemented using partially coherent illumination and standard optical components on an open-source microscope platform; together, these characteristics reduce acquisition burden and motion sensitivity, supporting dynamic imaging and resource-constrained deployment.

## 2. Theoretical Background

### 2.1. Gradient-Based Phase Imaging

Gradient-based phase imaging techniques recover optical phase by inverting a linearized intensity-to-phase transfer model under the WOA. Two widely used non-iterative approaches in this class are DPC [13, 16] and the TIE [21]. Both methods are compatible with partially coherent illumination and can be described within the weak object transfer function (WOTF) framework, but they exhibit fundamentally different spectral transfer characteristics.

DPC recovers phase gradients via asymmetric illumination and is characterized by an odd-symmetric transfer function, yielding a vanishing response as |**f**| → 0, where **f** = (*f*_*x*_, *f*_*y*_) denotes the two-dimensional spatial frequency. This odd symmetry gives directional sensitivity to phase gradients. In contrast, TIE relates axial intensity variations to the transverse Laplacian of phase. Under the uniform-intensity approximation, its Fourier-domain forward response scales as |**f**|^2^, while phase reconstruction requires an inverse-Laplacian operation with an ideal kernel proportional to 1/|**f**|^2^. Consequently, inversion must be regularized near zero spatial frequency.

Throughout this work, “low spatial frequency” refers to components approaching zero spatial frequency magnitude ( **f** 0), excluding the DC component (|**f**| →= 0), which corresponds to a global phase offset. This component is not recoverable in DPC due to the odd symmetry of its transfer function, while in TIE it is defined only up to an additive constant determined by boundary conditions.

### 2.2 Weak Object Approximation and NA Matching

In DPC microscopy, image formation is typically modeled under the WOA, where the sample transmission function

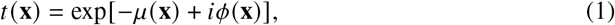

with **x** ∈ R^2^ denoting the transverse spatial coordinate, *µ*(**x**) the absorption (attenuation) distribution, and *ϕ*(**x**) the phase delay, is linearized via a first-order Taylor expansion as

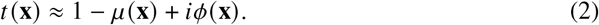

This approximation holds for weakly scattering or weakly absorbing specimens with small phase delays, typically |*ϕ*| ≲ 0.5 rad [22]. Accurate phase recovery in DPC further depends on the relationship between illumination and objective NAs, commonly parameterized by

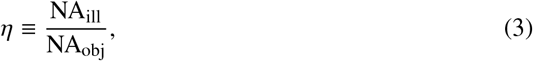

where NA_ill_ and NA_obj_ denote the illumination and objective NAs, respectively. When *η* ≈ 1, the illumination sufficiently fills the objective pupil, maximizing the spatial-frequency support of the phase transfer function (PTF) and reducing missing-frequency regions. This does not, however, eliminate the intrinsic low-frequency null of DPC. Operating at *η* < 1 improves model validity and robustness for thicker or higher-phase specimens, but reduces the support and magnitude of the PTF, limiting transfer efficiency and spatial resolution [19]. Conversely, increasing *η* increases susceptibility to WOA violations through higher-angle illumination and greater phase accumulation along oblique paths.

### 2.3 Polarization Differential Phase Contrast (pDPC)

pDPC [20, 23] is a snapshot implementation of DPC in which asymmetric illumination is encoded by polarization multiplexing and recovered in a single exposure using a division-of-focal-plane (DoFP) polarization sensor.

Let *I*_*θ*_ (**x**) denote the measured intensity at image-plane coordinate **x** = (*x, y*) corresponding to analyzer orientation *θ* ∈ {0°, 45°, 90°, 135°}. In the condenserless configuration, the polarization-structured illumination defines four effective source distributions associated with these analyzer channels. Synthetic asymmetric illumination pairs are formed as

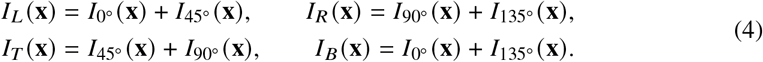

The normalized differential phase-contrast measurements are

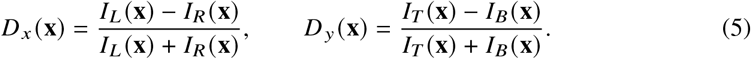

Polarization-resolved measurements are corrected by dark-background subtraction and application of a measured system-response matrix to compensate illumination imbalance and polarization-channel crosstalk [23]. The normalization in Eq. (5) further suppresses common-mode intensity variations and residual absorption contrast.

#### 2.3.1 WOTF-Based Forward Model

Using the linearized object model above, we adopt the Fourier-transform convention

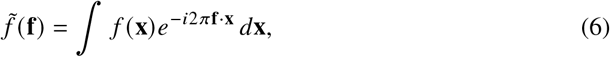

under which ∇^2^ ↔ −4π^2^|**f**|^2^.

Following the partially coherent image-formation model, the Fourier spectrum of each differential measurement is

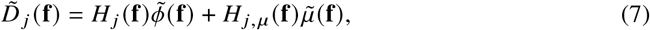

where *j* ∈ {*x, y*} indexes the effective illumination asymmetry, *H*_*j*_ (**f**) is the phase transfer function, and *H*_*j,µ*_ (**f**) describes absorption transfer.

The phase transfer function is obtained from the WOTF formalism [16, 24] as

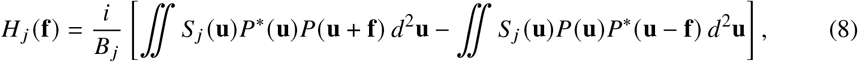

With

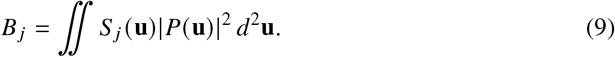

Here, **u** denotes the two-dimensional pupil-plane spatial-frequency coordinate, *S*_*j*_ (**u**) denotes the effective source distribution for the *j*-th asymmetric illumination pattern, *P* (**u**) is the objective pupil function, and *B*_*j*_ is the corresponding transmitted-intensity normalization factor. The resulting phase transfer function is predominantly imaginary and odd-symmetric, corresponding to sensitivity to phase gradients along the illumination-asymmetry axis.

Under the weak-absorption assumption, the absorption contribution enters primarily through the even-symmetric component of the transfer function and is partially suppressed by the differential measurements. Neglecting residual absorption–phase cross-talk, Eq. (7) therefore reduces to

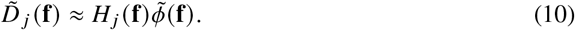

Since *D*_*j*_ (**x**) is real-valued, its Fourier transform satisfies Hermitian symmetry,

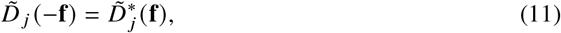

where ∗ denotes complex conjugation. The phase-only approximation assumes balanced illumination and negligible absorption–phase coupling.

#### 2.3.2. Phase Reconstruction

Semi-quantitative phase is recovered by solving the regularized least-squares inverse problem

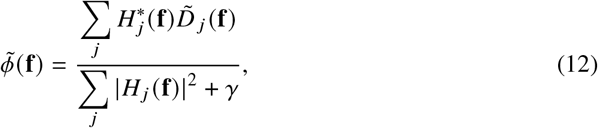

followed by inverse Fourier transformation. The Tikhonov parameter *γ* stabilizes the inversion near frequencies at which the combined transfer magnitude approaches zero. In practical pDPC implementations, deviations from ideal polarization response and illumination balance may introduce channel-dependent gain and cross-talk; these effects are corrected here using a measured system-response calibration [23].

### 2.4. Transport-of-Intensity Equation (TIE)

The TIE relates axial intensity variations to transverse phase gradients. The governing equation is

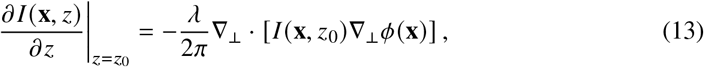

where λ is the illumination wavelength and ∇_⊥_ denotes the transverse gradient operator. For approximately uniform in-focus intensity, *I* (**x**, *z*_0_) ≈ *I*_0_, Eq. (13) reduces to

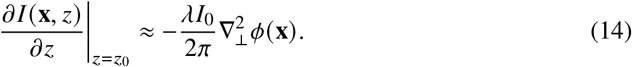

Using the Fourier-transform convention defined above,

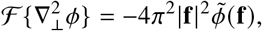

the corresponding ideal inverse-Laplacian solution is

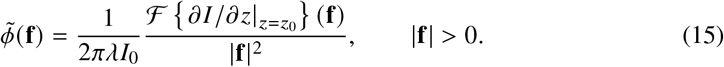

#### 2.4.1. Phase Reconstruction

For spatially non-uniform in-focus intensity, phase can be recovered using the FFT-based formulation [25]. Defining

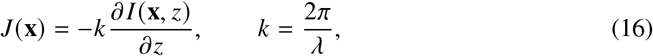

the reconstruction follows Teague’s auxiliary-function formulation [21], which assumes that the intensity-weighted phase-gradient field is irrotational and may be written as *I* (**x**, *z*_0_)∇_⊥_*ϕ*(**x**) = ∇_⊥_*Ψ*(**x**), where *Ψ* is an auxiliary potential. Under this assumption,

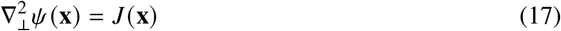

is solved using a Tikhonov-regularized inverse Laplacian. The phase gradient is then evaluated as

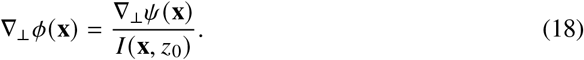

Taking the divergence gives

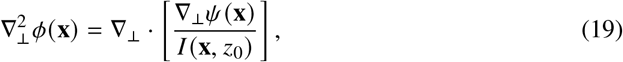

from which *ϕ* (**x**) is recovered using a second Tikhonov-regularized inverse Laplacian. The regularization stabilizes the inversion near zero spatial frequency.

## 3. Methods

### 3.1. System Overview

All experiments were conducted on an *openFrame* modular open-source inverted microscope platform [26], customized to support reported polarization-encoded pDPC imaging [20, 23, 27] and TIE phase recovery within a single optical setup (Fig. S1). A polarization-sensitive DoFP camera (Blackfly BFS-U3-51S5P-C, Teledyne FLIR) simultaneously recorded four linear polarization channels. The optical train (Fig. 1(b)) comprised a transillumination subsystem (unified hpDPC illumination module, (Fig. 1(a))), sample plane, objective lens, tube lens, and polarization-sensitive camera. The unified hpDPC illumination module was custom designed to provide frequency-selective dual-modality illumination for pDPC and TIE without mechanical reconfiguration.

**Fig. 1.**
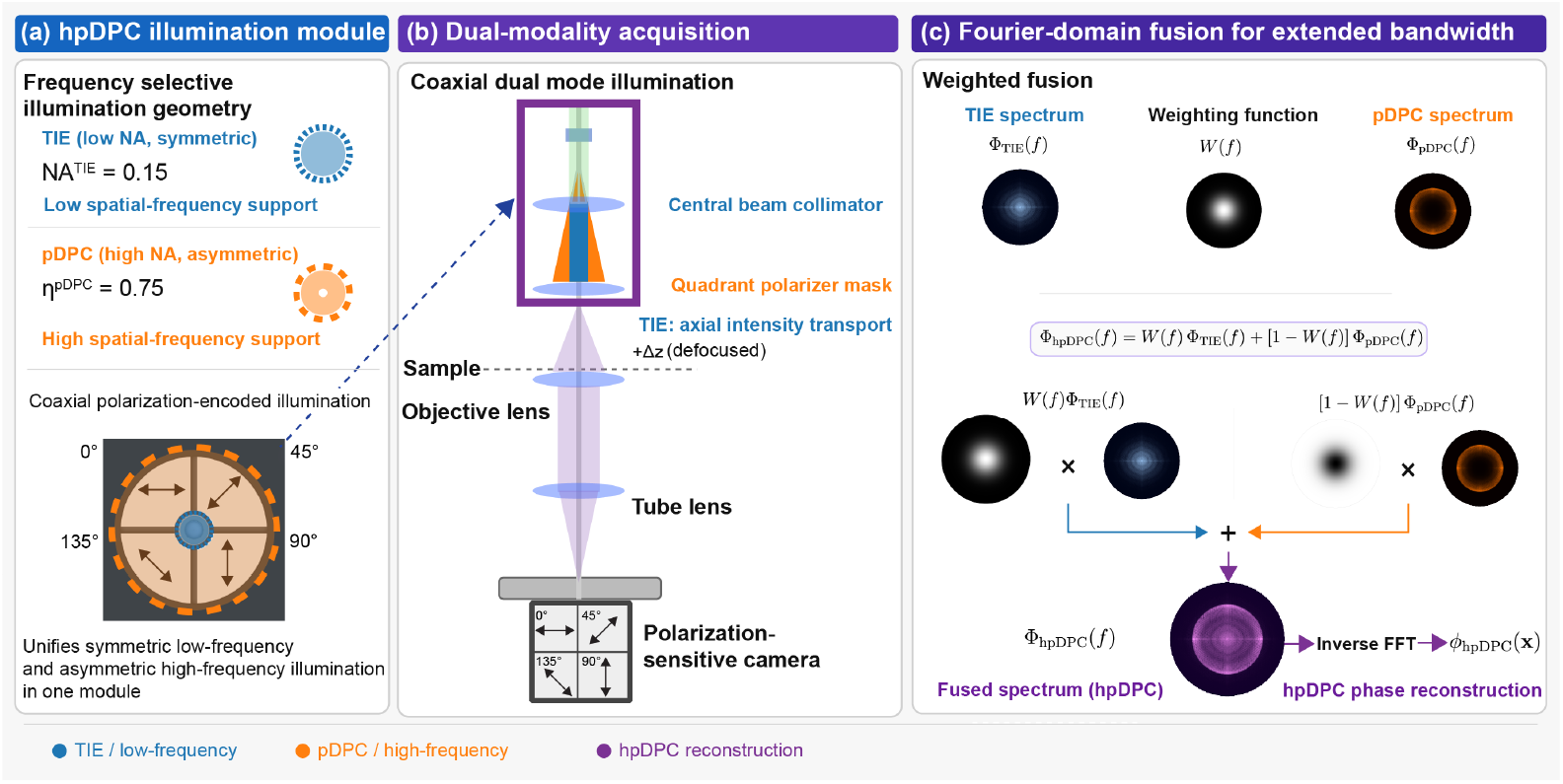
Hybrid polarization differential phase contrast (hpDPC) imaging architecture. (a) Unified illumination module combining low-NA symmetric illumination for TIE with high-NA polarization-encoded asymmetric illumination for pDPC using a quadrant polarizer mask. (b) Snapshot pDPC and defocused TIE measurements are acquired via a polarization-sensitive camera and axial defocus (Δ*z*). (c) Complementary Fourier-domain transfer characteristics of pDPC (high spatial frequency) and TIE (low spatial frequency) enable frequency-selective fusion for extended spatial-frequency coverage.

### 3.2. Unified hpDPC Illumination Module

The unified illumination module (Fig. 1(a)) coaxially integrates polarization-encoded asymmetric illumination for pDPC with symmetric illumination for TIE in a compact assembly. Unlike quadrant polarizer masks optimized solely for pDPC, the design partitions illumination angles between the two phase-recovery modalities. A condenserless architecture was adopted so that the illumination angles are defined by source geometry rather than a condenser lens, thereby providing angular separation between the pDPC and TIE channels. Although this departs from Köhler illumination, the effective pupil function remains well defined under geometric optics, allowing the linearized DPC forward model under the WOA to be applied.

The pDPC path employs a four-quadrant linear polarization assembly (0°, 45°, 90°, and 135°) constructed from cut linear polarizing film (XP42-40, Edmund Optics), with tracing paper positioned behind the film to provide diffusive illumination (Fig. 1(b)). In combination with the DoFP camera, this produces four asymmetric illumination conditions in a single exposure, enabling snapshot recovery of two-dimensional phase gradients without temporal multiplexing and supporting dynamic or live-cell imaging. Illumination is provided by three high-power LEDs (LEDENGIN LZ1-10G102, OSRAM; peak wavelength 523 nm), arranged radially around the central TIE channel and mounted in a custom 3D-printed cradle with a shared heatsink. The LEDs are synchronized to the camera exposure to provide high photon flux during acquisition while reducing the average photon dose during live-cell or time-lapse imaging. The central inactive aperture produces a low-angle obscuration that suppresses low-frequency contributions to the pDPC transfer function while preserving high-angle asymmetric illumination. Although the obscuration reduces transmitted power, exposure times were adjusted to maintain comparable photon counts between modalities. The effective pDPC illumination NA is determined geometrically by the lateral displacement of the LEDs and polarization mask relative to the optical axis and their distance from the sample plane; the transillumination-pillar height was adjusted for each objective to maintain *η*_pDPC_ ≈ 0.75.

The coaxial TIE channel uses a fiber-coupled LED (LEDENGIN LZ1-10G102, OSRAM) with a 105 µm core-diameter fiber to provide low-angle symmetric illumination. The output is collimated using a fiber collimator (F280APC-A, Thorlabs) and further constrained by a cylindrical aperture (12 mm diameter, 40 mm length), yielding an effective illumination NA of approximately 0.15. This low NA provides sensitivity to low-spatial-frequency phase while supporting stable TIE inversion. Focus–defocus displacement of the sample is performed using an electronic stage (Nanomax 381, Thorlabs).

All illumination sources are driven by custom current-controlled drivers and synchronized with the camera exposure using an Arduino Nano microcontroller, providing precise timing between pDPC and TIE acquisitions while maximizing photon flux and minimizing thermal load.

### 3.3. Data Acquisition Protocol

Microscope control and image acquisition were automated via a custom software routine. A single hpDPC acquisition cycle comprises three measurements (Fig. 2(a)): one polarization-multiplexed pDPC frame and two intensity images for TIE, acquired at the nominal focal plane *z*_0_ and at a single defocused position (*z*_0_ + Δ*z*). The defocused image was acquired following a controlled axial displacement of the sample by Δ*z*, after which the system was returned to the nominal focal plane. This protocol reduces acquisition count compared to existing multi-shot DPC–TIE approaches [19]. The defocus distance Δ*z* was predefined according to the sample and imaging conditions, typically within 0.5–10 µm in this study, while maintaining the finite-difference approximation within the paraxial TIE regime [25].

**Fig. 2.**
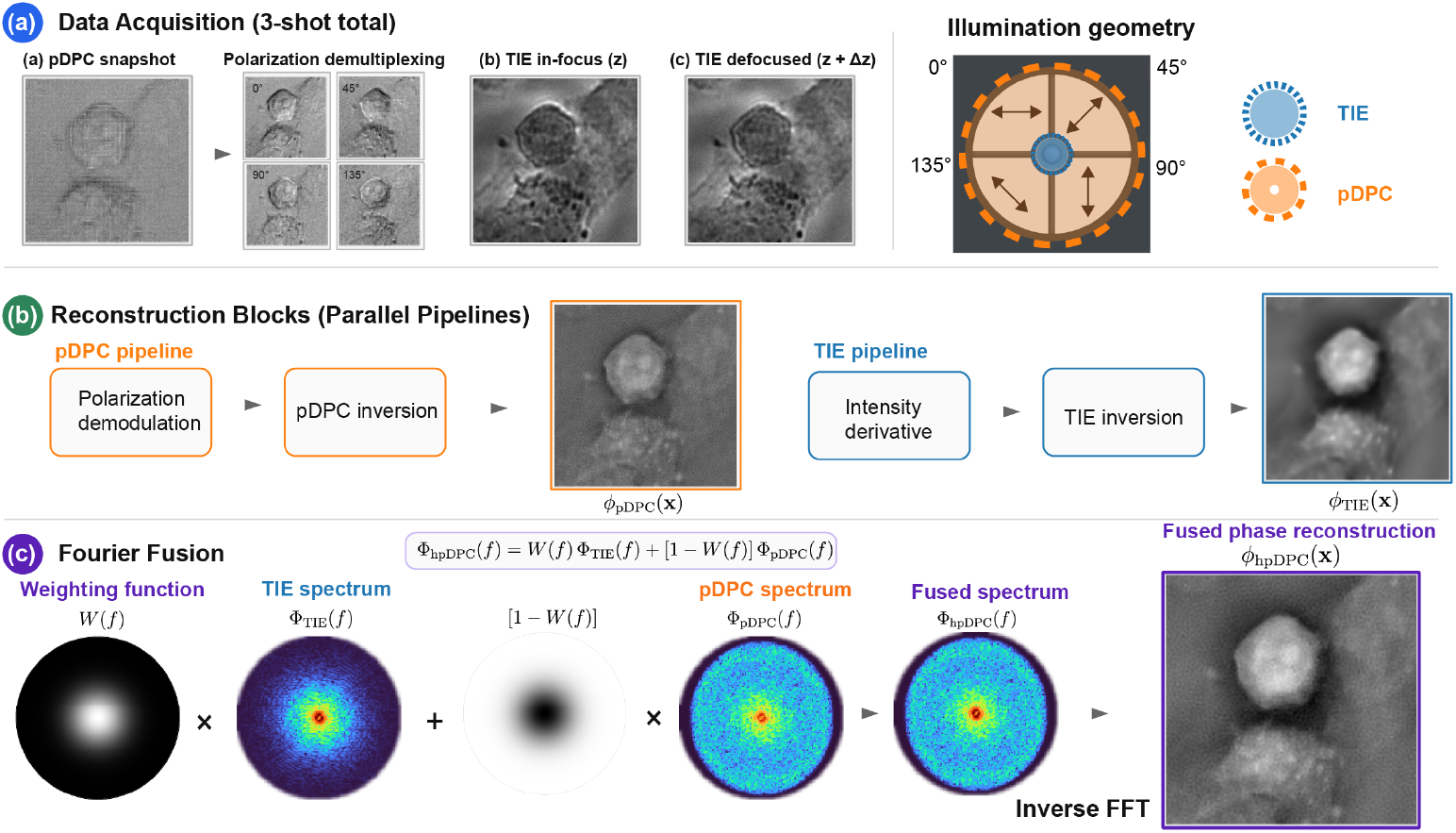
Overview of the hpDPC reconstruction and fusion pipeline. (a) Data acquisition: A polarization-multiplexed pDPC image is demultiplexed into four analyzer orientations, while in-focus and defocused intensity images are acquired for TIE. (b) Reconstruction: Parallel pipelines recover phase, with pDPC capturing high-spatial-frequency content via transfer function inversion and TIE recovering low-spatial-frequency content via the inverse Laplacian. (c) Fourier-domain fusion: The phase estimates are combined using a frequency-dependent weighting function 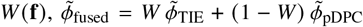, followed by inverse Fourier transformation to obtain the final phase map.

### 3.4. Reconstruction Implementation

For pDPC reconstruction (Fig. 2(b)), following dark-background subtraction and system-response calibration, the polarization-sensitive image was demultiplexed into the four analyzer channels (0°, 45°, 90°, and 135°), which were combined to form the left–right and top–bottom asymmetric illumination images. The normalized differential measurements were computed using Eq. (5), and phase was reconstructed using the regularized WOTF inversion of Eq. (12).

For TIE reconstruction (Fig. 2(b)), an in-focus image *I* (**x**, *z*_0_) and a single defocused image *I* (**x**, *z*_0_ + Δ*z*) were acquired under symmetric low-NA illumination. The axial intensity derivative was approximated using the forward finite difference

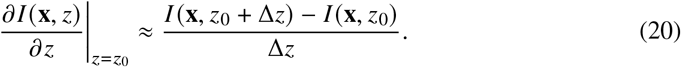

Phase was then reconstructed using the non-uniform-intensity formulation of Eqs. (16)–(19). A lower intensity threshold was imposed on *I* (**x**, *z*_0_) before the division in Eq. (18) to prevent numerical instability in low-intensity regions. The reconstruction parameters used throughout the experiments are summarized in Table S1 of Supplement 1.

### Spectral Fusion of pDPC and TIE

Based on their differing spectral responses, the independently reconstructed pDPC and TIE phase maps were combined in the Fourier domain, with TIE weighted toward low spatial frequencies and pDPC toward higher spatial frequencies. Prior to fusion, the pDPC reconstruction, obtained on the reduced sampling grid associated with polarization-channel demosaicing, was upsampled by a factor of two to match the TIE sampling grid. This establishes a common computational grid for Fourier-domain fusion but does not extend the native spatial-frequency support of the pDPC reconstruction.

The two reconstructions were subsequently combined in the Fourier domain using complementary radial weights (Fig. 2(c)). Let 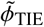(**f**) and 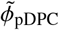(**f**) denote the Fourier transforms of the TIE and upsampled pDPC phase reconstructions, respectively. The fused spectrum is

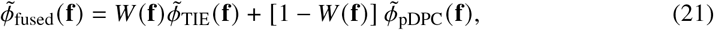

where the normalized radial spatial frequency is

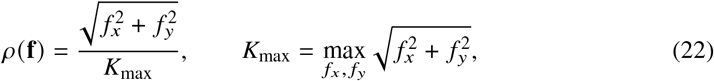

and the radially symmetric TIE weighting function is

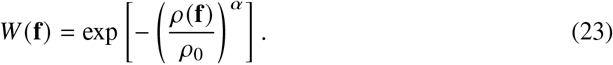

For all reconstructions, *ρ*_0_ = 0.05 and *α* = 2, where *α* is a weighting roll-off exponent. Since *W* (**0**) = 1, the zero-frequency term is assigned to the TIE reconstruction, while the pDPC contribution increases smoothly with radial spatial frequency. Because *ρ* is normalized by *K*_max_, *ρ*_0_ is a dimensionless empirical transition parameter rather than an NA-defined cutoff; its corresponding physical transition frequency scales with *K*_max_ and therefore with the spatial sampling. No inter-modality gain or amplitude normalization was applied prior to fusion.

Following spectral combination, a weak frequency-dependent regularization was applied,

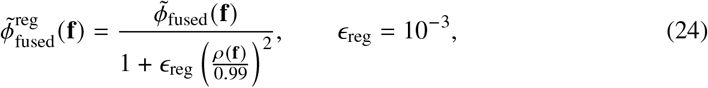

after which the fused phase was obtained by inverse Fourier transformation,

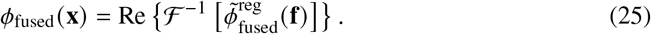

Because the global phase offset is arbitrary, the spatial mean of the fused reconstruction was subsequently matched to that of the TIE reconstruction,

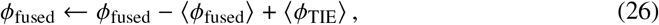

where ⟨·⟩ denotes the spatial mean over the reconstruction domain.

The fusion is empirical and does not explicitly incorporate the individual phase transfer functions or noise power spectra of the two modalities. Instead, their relative contributions are prescribed by the smooth radial weighting in Eq. (23).

The hpDPC spectral-fusion and analysis routines were implemented in MATLAB® R2024b and Python 3.9.7. Code for fusing precomputed pDPC and TIE phase reconstructions and reproducing the associated analyses is available in the repository specified in the Data Availability statement.

### 3.6. Sample Preparation

#### 3.6.1. Preparation of Microspheres

For preparation of polystyrene (PS) microspheres, 5 µl of 4 µm diameter spheres from the Flow Cytometry Size Calibration Kit (F-13838, Thermo Fisher Scientific) were spotted onto poly-l-lysine (P4707, Merck)-coated coverslips and air dried before mounting with ProLong Glass Antifade Mountant (P36980, Thermo Fisher Scientific) according to the manufacturer’s instructions.

#### 3.6.2. Preparation of A549 Cells

A549 cells (human lung adenocarcinoma epithelial cell line) were cultured in Dulbecco’s Modified Eagle Medium (DMEM) supplemented with 10% fetal bovine serum (FBS) and 1% penicillin–streptomycin at 37 °C and 5% CO_2_ in a humidified atmosphere. For seeding onto coverslips, cells were trypsinized and seeded at a density of 1 × 104 cells cm−2. Following overnight incubation, A549 cells were washed three times with 1 × phosphate buffered saline (PBS) (pH 7.4) and fixed with 4% paraformaldehyde (PFA) in PBS for 15 min at 37 °C before a final three washes with PBS. Coverslips were mounted onto slides with PBS and sealed with nail varnish.

#### 3.6.3. Preparation of Mitomycin C-treated *M. smegmatis*

The ΔimuB::mEos4a-ImuB reporter strain, generated from an ImuB-deficient MC2155 *M. smegmatis* strain [28] electroporated with an integrative pAINT plasmid, was cultured in Middlebrook 7H9 broth supplemented with 0.2% glycerol, 0.05% Tween-80, and 10% (oleic acid-albumin-dextrose-catalase) OADC at 37 °C with shaking. DNA damage and cell elongation were induced by addition of 0.2 µg/ml Mitomycin C at OD_600_ 0.2. Following 4 h of growth at 37 °C with shaking, the sample was immobilized on a 1% (w/v) agarose pad. Briefly, the agarose pad was prepared by casting 1% low-gelling-temperature agarose (A9414, Merck) between two glass slides. Following solidification of the agarose and removal of one slide, 3–6 µl of bacterial suspension was deposited onto the agarose and allowed to dry for 1–5 min to facilitate cell adhesion before application of the coverslip and imaging.

#### 3.6.4. Preparation of Heat-Treated and Control Samples for Learning-Based Cell State from Phase

*M. smegmatis* MC2155 was cultured in Middlebrook 7H9 broth supplemented with 0.2% glycerol, 0.05% Tween-80, and 10% OADC at 37 °C with shaking. Mid-to late-log-phase cultures (OD_600_ 0.3–1.0) were either heated at 65 °C for 15 min (heat-treated) or maintained at 37 °C (control). Thereafter, control and heat-treated samples were pelleted by centrifugation at 3000×*g* at room temperature (RT) for 3 min, resuspended in 0.1 µM SYTOX Orange Dead Cell Stain (S34861, Thermo Fisher Scientific) in 1× phosphate-buffered saline (PBS; pH 7.4) (10016-023, Gibco), and incubated for 15 min at RT to label cells with compromised membrane integrity. Following incubation, cells were passed through a 5 µm filter to minimize cell clumping before being deposited onto a 1% agarose pad as described above.

## 4. Simulations

Simulations were performed to evaluate the spectral behavior of pDPC and TIE and to assess hpDPC under departures from the WOA. Synthetic measurements were generated using a partially coherent forward model with the full pure-phase transmittance *t* (**x**) = exp[*iϕ* **x**], corresponding to *µ* (**x**) = 0 in Eq. 1. For each discrete illumination source point, the object spectrum was shifted, pupil-filtered, and propagated to the image plane, and the resulting intensities were incoherently summed. Defocused TIE measurements were generated analogously after propagation to *z* = *z*_0_ + Δ*z*. The simulated data were reconstructed using regularized WOTF-based pDPC inversion and the TIE method described above, thereby retaining nonlinear phase effects in the forward model that are absent from the WOA-based pDPC reconstruction.

### 4.1. UCT Logo and Microlens Study

All restricted pDPC simulations use an illumination–objective mismatch parameter of *η*_pDPC_ = 0.75, consistent with regimes reported to improve robustness to WOA violations [19] while retaining useful high-spatial-frequency transfer; the corresponding reduction in low-spatial-frequency sensitivity is compensated by TIE. Additionally, an NA-matched pDPC configuration 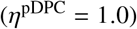 is included for reference. For TIE, the illumination NA is fixed at 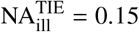, providing low-frequency sensitivity while maintaining numerical stability of the inverse Laplacian and support under moderate WOA violations. For conventional circular illumination, increasing 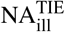 does not improve high-frequency phase recovery but rather reduces phase contrast as the transfer function weakens and vanishes near NA_ill_ → NA_obj_, leading to poor SNR and loss of recoverable phase information [29]. A fixed 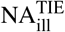 ensures consistent low-frequency coverage across objectives, reducing the need for illumination reconfiguration.

To evaluate the expected behavior of pDPC and TIE and validate the proposed hpDPC framework, simulations were performed using a phase object containing both low- and high-spatial-frequency content (Fig. 3(a)). A microlens phase profile with embossed text was used as ground truth phase object, combining smoothly varying lens curvature with sharp character phase gradients to challenge phase reconstruction across spatial frequencies.

**Fig. 3.**
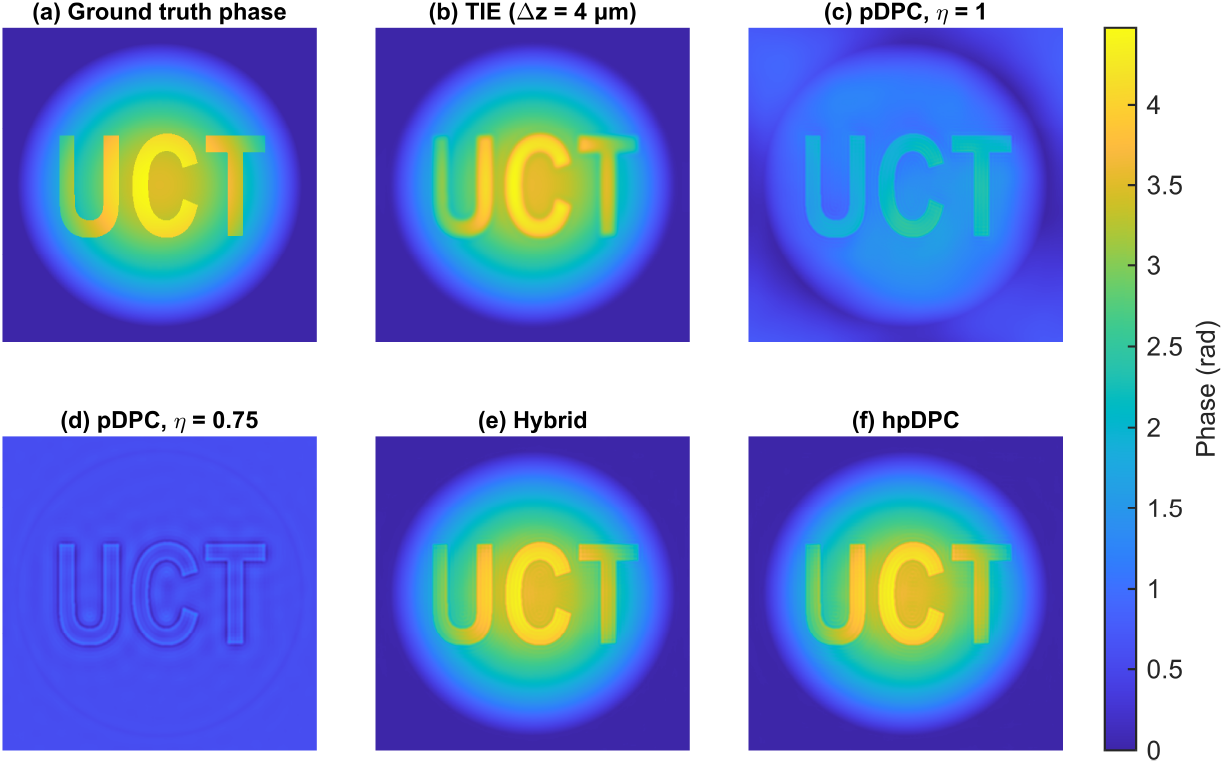
Simulation results comparing phase retrieval methods. (a) Ground-truth phase profile of a microlens array (3.5 rad height) superposed with high-frequency text (“UCT”). (b) TIE reconstruction (Δ*z* = 4 µm). (c) pDPC reconstruction (matched illumination, *η* = 1). (d) pDPC reconstruction (unmatched illumination, *η* = 0.75) of the same array. (e) Hybrid TIE and pDPC reconstruction without central obscuration in the illumination aperture. (f) Proposed hpDPC reconstruction using annular illumination with central obscuration (NA_hole_ ≈0.15), analogous to that of a combined modality illumination module. Phase in radians.

#### 4.1.1. Baseline Simulations

We compared TIE, pDPC with matched and unmatched illumination NA, and hybrid TIE–pDPC reconstructions with and without a central obscuration (Fig. 3(b–f)). This progression isolates the individual modality limitations and the effect of the proposed illumination geometry.

For TIE (Fig. 3(b)), symmetric illumination with Δ*z* = 4 µm was used to balance numerical stability and low-spatial-frequency sensitivity. For matched pDPC (Fig. 3(c)), quadrant asymmetric illumination without a central obscuration was used with NA_ill_ ≈NA_obj_, providing near-optimal PTF support under the WOA. TIE recovers the slowly varying microlens profile but attenuates high-frequency edges, whereas pDPC preserves the fine text features but underestimates the low-frequency phase background. Reducing the pDPC illumination NA further increases this low-frequency underestimation (Fig. 3(d)).

The hybrid reconstruction without central obscuration (Fig. 3(e)) combines TIE with unmatched pDPC and recovers both the slowly varying lens profile and sharp phase features, despite the WOA violation. In hpDPC (Fig. 3(f)), the unmatched pDPC illumination additionally incorporates the central obscuration required by the coaxial TIE channel. The resulting reconstruction retains both large-scale and fine structural features while reproducing the illumination geometry used experimentally. These results demonstrate the complementary spectral response of TIE and pDPC and motivate the frequency-selective fusion used in hpDPC.

#### 4.1.2. Role of the Central Obscuration

The pDPC PTF depends strongly on the illumination distribution, with illumination matched to the objective NA providing the broadest and most uniform spatial-frequency response [30]. To determine how the central obscuration used in hpDPC modifies this response, we compared the pDPC component at *η* = 0.75 with and without suppression of low-angle illumination. The central obscuration preferentially attenuates the pDPC response near zero spatial frequency while largely preserving higher-frequency phase-gradient transfer. This is evident from the difference map, radially averaged spectra, and extracted phase profiles in Fig. S2(c–e), which show reduced slowly varying phase content with minimal loss of fine structural detail.

To quantify reconstruction performance, relative spectral error was computed from the Fourier-domain reconstruction error normalized by the ground-truth spectral power and radially averaged over annular spatial-frequency bins. Root-mean-square-error (RMSE) was calculated over the full simulation domain, while structural similarity index measure (SSIM) was evaluated after jointly normalizing each reconstruction and the ground truth to [0, 1]. Fig. 4 shows that TIE exhibits greater relative spectral error at high spatial frequencies, whereas pDPC exhibits greater low-frequency error (Fig. 4(a,b)). Both hybrid reconstructions reduce error across these regimes, with hpDPC retaining high-frequency recovery despite the central obscuration. The RMSE and SSIM metrics in Fig. 4(c) support the same trend, with the hybrid and hpDPC methods achieving low RMSE and high SSIM.

**Fig. 4.**
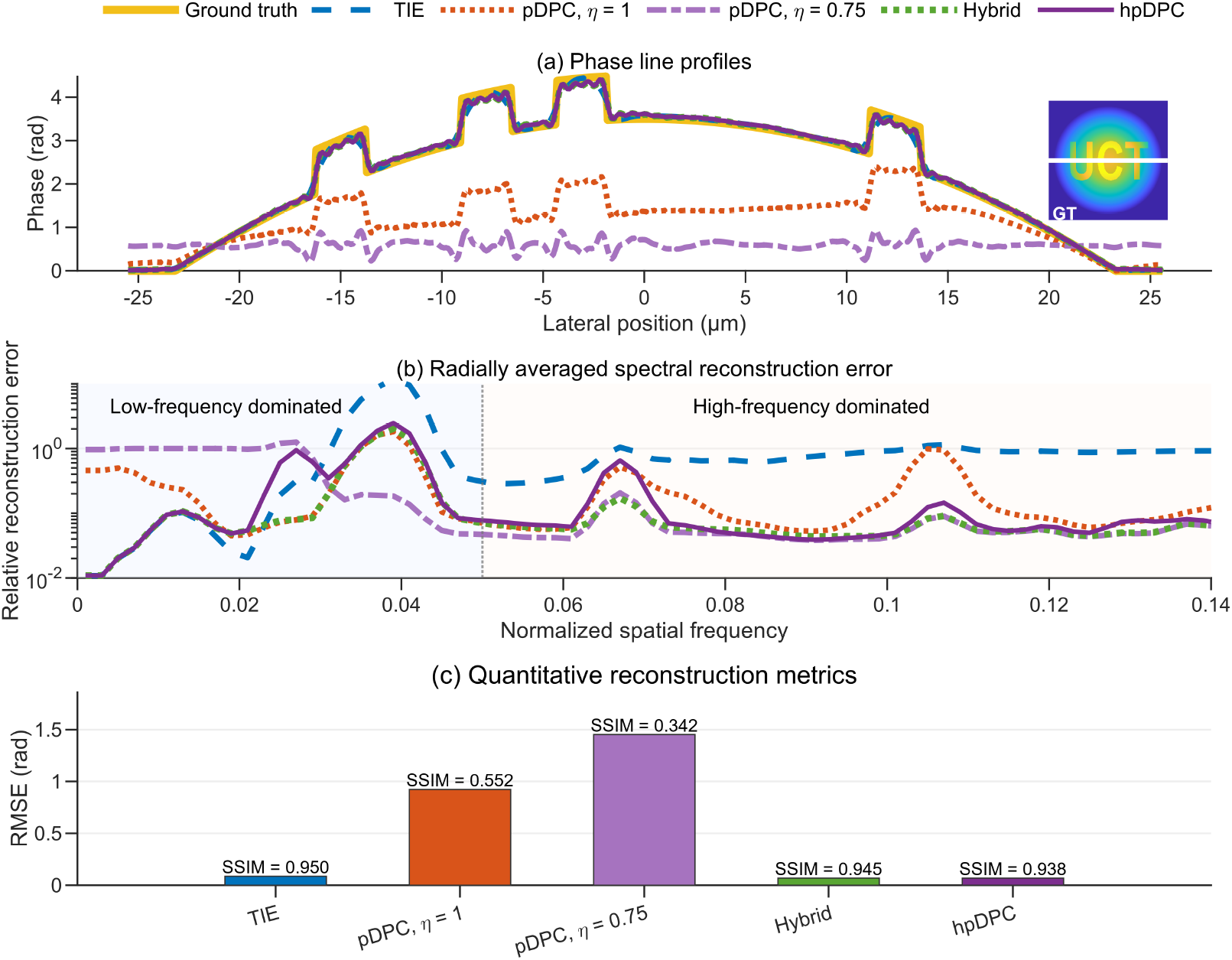
Quantitative comparison of phase reconstruction performance across spatial frequencies, comparing TIE, pDPC with matched-and unmatched NA, and hybrid TIE and pDPC, with and without a central obscuration. (a) Phase reconstruction line profiles through the simulated microlens object containing low-frequency curvature and high-frequency text features (“UCT”). Inset: ground-truth phase object and extracted profile location. (b) Radially averaged spectral reconstruction error versus normalized spatial frequency. (c) Quantitative reconstruction metrics. Bars indicate root-mean-square-error (RMSE) relative to the ground truth, with annotated structural similarity index measure (SSIM) values.

## 5. Experiments and Results

### 5.1. Microspheres

A sample of 4 µm PS microspheres (*n*_*b*_ = 1.603 near λ = 523 nm) mounted in ProLong Glass Antifade Mountant (*n*_*m*_ = 1.52, at 589 nm) was used to evaluate the phase response of the three reconstruction methods with 20 × 0.4 NA and 40 × 0.75 NA objective lenses, where *n* denotes the RI. For TIE reconstruction, intensity images were acquired at the object plane and at a defocus position of Δ*z* = 5 µm.

For a homogeneous sphere under ideal embedding conditions, the maximum phase shift for central traversal is

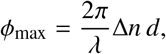

where Δ*n* = *n*_*b*_ − *n*_*m*_ and *d* = 4 µm. The quoted mountant RI is specified at the sodium D-line (589 nm), and wavelength-resolved dispersion data for the cured medium are unavailable. Using the reported PS value *n*_*b*_ ≈ 1.603 near the experimental wavelength and *n*_*m*_ ≈ 1.52 gives Δ*n* ≈ 0.083 and, at λ = 523 nm, *ϕ*_max_ ≈ 3.99 rad.

The corresponding phase maps and line profiles are shown in Fig. 5. The profile in Fig. 5(b), extracted along the red line in Fig. 5(a), intersects three microspheres, passing approximately through the centre of the middle sphere and the peripheral regions of the outer two. The latter therefore have shorter optical traversal lengths, scaling as 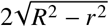 for lateral offset *r* from the sphere centre, where *R* = *d*/2.

**Fig. 5.**
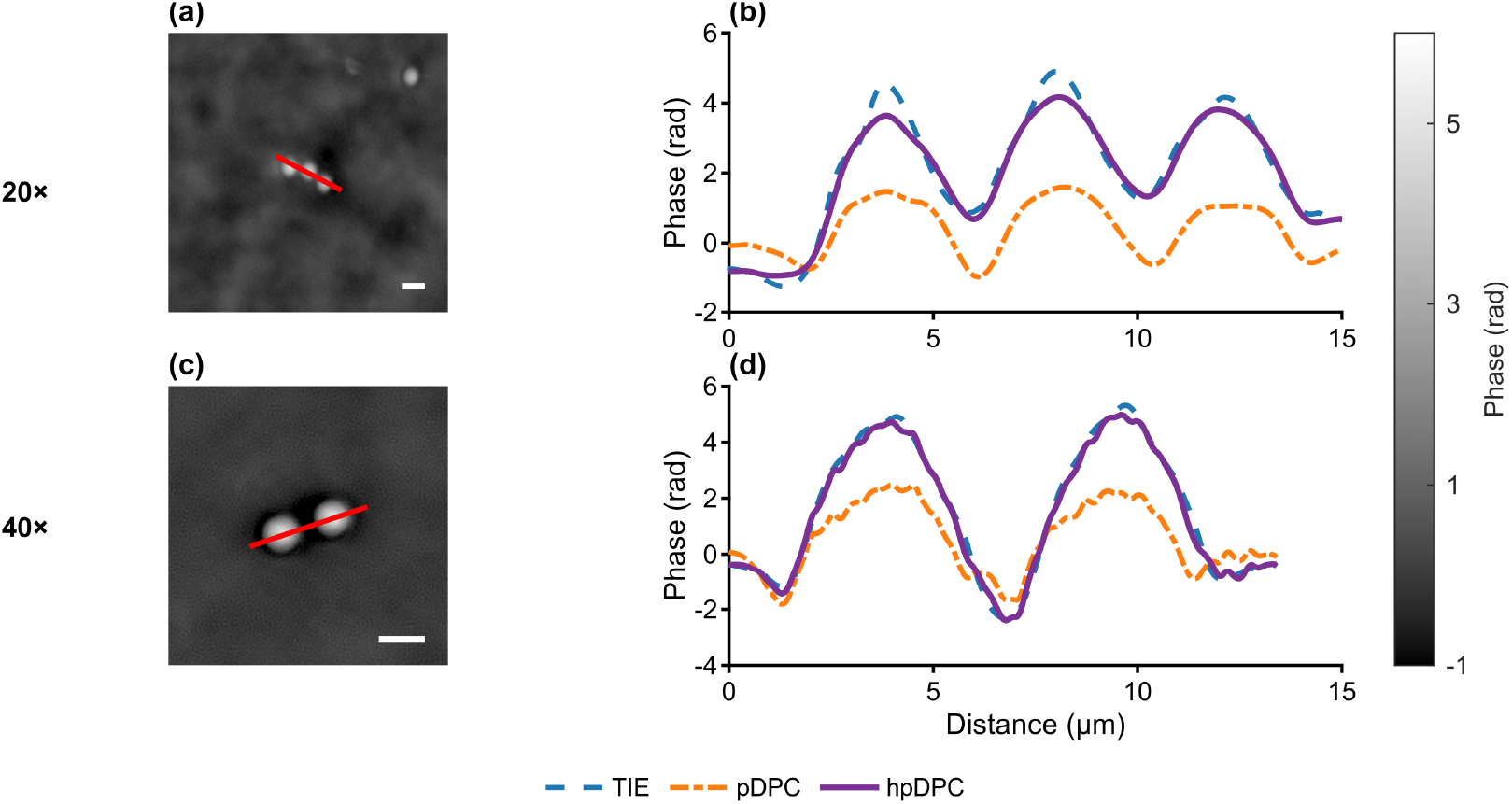
Phase validation using 4 µm polystyrene (PS) microspheres (*n*_*b*_ = 1.603, *n*_*m*_ = 1.52). (a,c) hpDPC phase reconstructions at 20 × and 40 × magnification, respectively, with red lines indicating the extracted phase profiles. Scale bars: 5 µm. (b,d) Corresponding phase line profiles comparing TIE, pDPC, and hpDPC reconstructions. TIE recovers peak phase values near the theoretical maximum phase shift (*ϕ*_max_ ≈ 3.99 rad), while pDPC underestimates the phase amplitude due to reduced low-spatial-frequency sensitivity. The proposed hpDPC reconstruction improves phase recovery and overall agreement with the expected phase response. Phase in radians.

The TIE and hpDPC profiles in Fig. 5(b) follow the expected geometry, with lower phase amplitudes for the outer spheres due to their shorter traversal lengths. pDPC systematically underestimates phase amplitude and exhibits reduced dynamic range, consistent with its lower sensitivity to slowly varying phase. hpDPC closely follows TIE in both phase magnitude and profile morphology.

Fig. 5(c,d) shows the corresponding hpDPC reconstruction and line profiles for a spaced microsphere pair at 40 ×. Relative to the 20 × results, the higher-NA reconstruction exhibits sharper boundaries and finer spatial detail. The reconstructed peak phase exceeds the theoretical prediction and weak negative edge lobes are present, consistent with transfer-function inversion artefacts. The amplitude discrepancy may additionally reflect uncertainty in the cured-mountant RI at 523 nm, microsphere diameter and RI tolerances, and non-ideal embedding following air drying on the poly-l-lysine-coated coverslip. Despite these uncertainties, hpDPC improves phase recovery relative to pDPC while remaining in close agreement with TIE in phase morphology and amplitude.

### 5.2. A549 Cells

PFA-fixed A549 cells were imaged with a 20 × 0.4 NA objective lens. For TIE reconstruction, intensity images were acquired at the object plane and at a defocus position Δ*z* = 5 µm. Fig. 6(a) and (b) compare phase reconstructions obtained using pDPC and TIE, respectively. The pDPC reconstruction emphasizes fine structural detail and sharp cellular boundaries, but exhibits reduced sensitivity to slowly varying phase within the cell interior. In contrast, the TIE reconstruction recovers smooth and physically consistent phase across the cell body, with diminished edge definition due to attenuation of fine-scale features. Fig. 6(c) presents the hpDPC reconstruction that preserves interior phase consistency while restoring structural detail at cellular boundaries, yielding a more complete representation of the sample. An enlarged comparison in Fig. 6(d–f) further highlights this behavior.

**Fig. 6.**
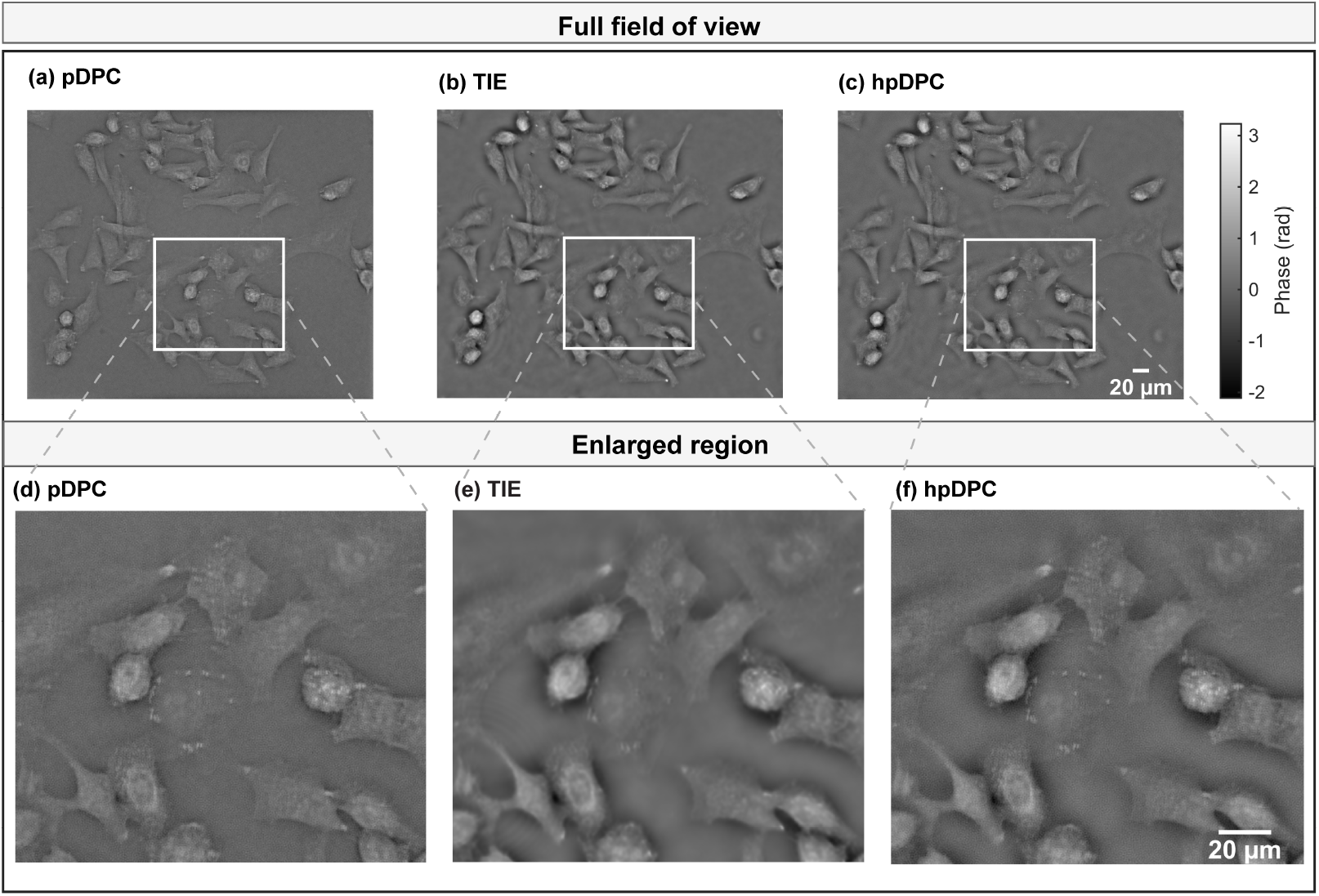
Comparison of phase reconstructions for A549 cells using (a, d) pDPC, (b, e) TIE, and (c, f) hpDPC. Top row: full field of view. Bottom row: magnified regions corresponding to the boxed ROI. Scale bars: 20 µm. Phase in radians.

### 5.3. M. smegmatis (60×)

Phase reconstructions of *M. smegmatis* imaged with a 60 × 0.8 NA objective lens are shown in Fig. 7. For TIE reconstruction, intensity images were acquired at the object plane and at a defocus position of Δ*z* = 5 µm. The full-field phase reconstructions in Fig. 7(a–c) illustrate differences in spatial-frequency response and phase fidelity across methods. pDPC preserves fine structural detail but under-recovers phase amplitude, impacting contrast. In contrast, the TIE reconstruction retains stronger low-frequency phase information at the expense of high-frequency attenuation, resulting in blurred features, ringing artifacts, and reduced separability of adjacent bacilli. The proposed hpDPC method balances these effects by recovering higher phase amplitude while preserving structural sharpness, thereby improving contrast and feature definition. The enlarged regions in Fig. 7(d–f) further highlight these trends. pDPC maintains edge sharpness but compresses peak phase, whereas TIE exhibits pronounced low-pass behavior and loss of fine structural detail. hpDPC mitigates both limitations, producing sharper feature boundaries with improved agreement with the expected phase structure of the bacilli.

**Fig. 7.**
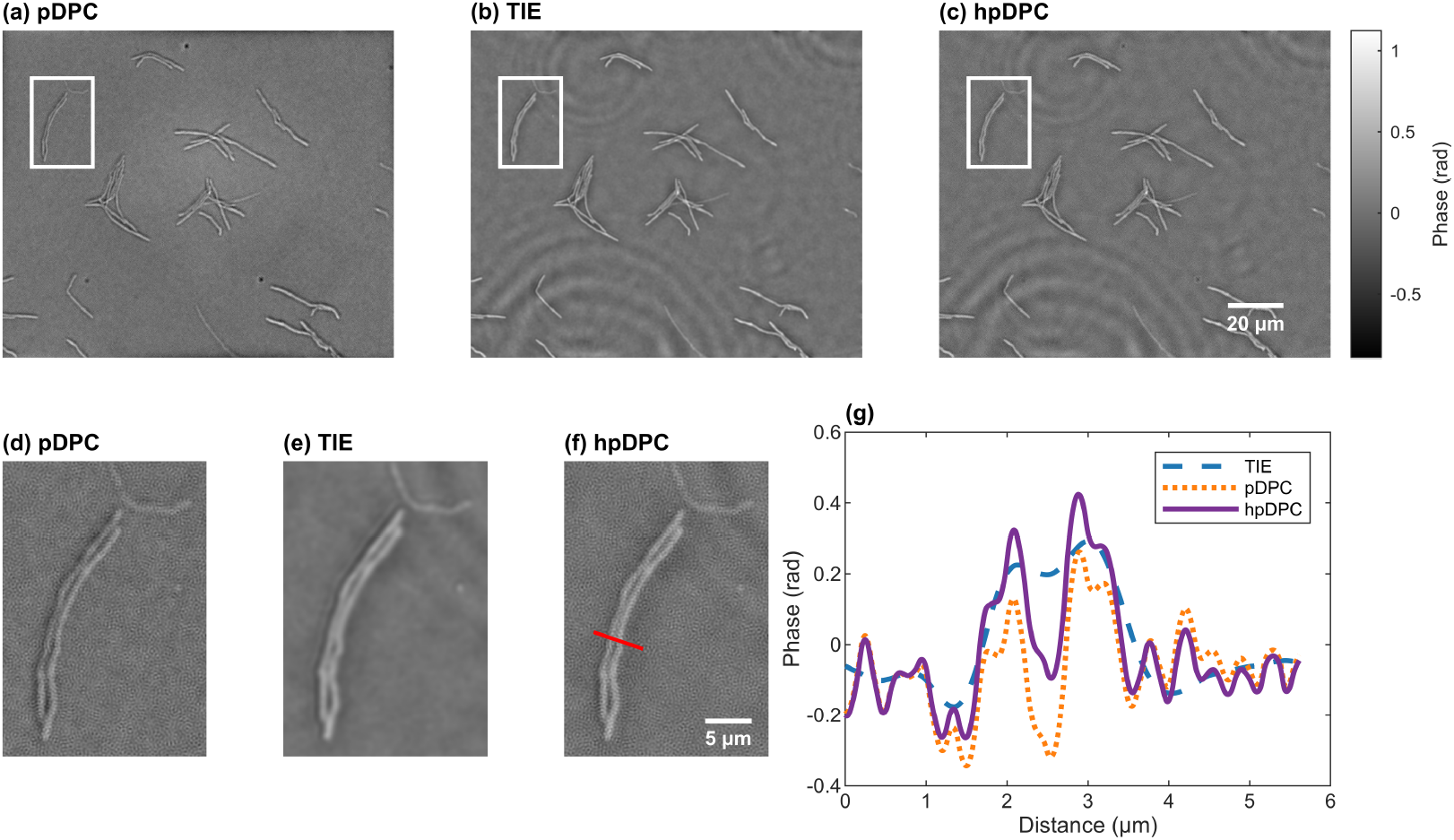
Comparison of phase reconstructions of representative *M. smegmatis* bacilli at 60 × using pDPC, TIE, and hpDPC. (a–c) Full field-of-view phase reconstructions obtained using pDPC, TIE, and hpDPC, respectively. (d–f) Magnified views corresponding to the boxed regions in (a–c). The red line in (f) indicates the location of the extracted phase profiles shown in (g), comparing TIE, pDPC, and hpDPC reconstructions. Scale bars: 20 µm in (c) and 5 µm in (f). Phase in radians.

A quantitative comparison is shown via line profiles in Fig. 7(g), extracted along the transect indicated in Fig. 7(f). The transect intersects two adjacent bacilli, giving an expected two-peak phase profile. TIE fails to resolve this structure, yielding a single broadened peak, whereas pDPC resolves the two peaks, consistent with improved high-spatial-frequency recovery. hpDPC preserves the two-peak morphology while improving the recovered phase profile.

### 5.4. M. smegmatis (40×)

*M. smegmatis* samples were additionally imaged with a 40 × 0.75 NA objective lens to assess performance at reduced spatial resolution (Fig. 8). Full-field reconstructions reflect the expected frequency responses. pDPC (Fig. 8(a)) preserves fine structural detail but underestimates phase amplitude. TIE (Fig. 8(b)) emphasizes lower spatial frequencies, producing smoother but blurred phase maps with diminished visibility of fine features and pronounced low-frequency background artifacts. The proposed hpDPC reconstruction (Fig. 8(c)) combines these characteristics, preserving structural detail while restoring phase contrast. Magnified views (Fig. 8(d–f)) reinforce these trends. pDPC resolves fine structure, whereas TIE fails to separate closely spaced bacilli. hpDPC preserves the edge sharpness of pDPC while improving feature separability relative to TIE and restoring phase contrast relative to pDPC.

**Fig. 8.**
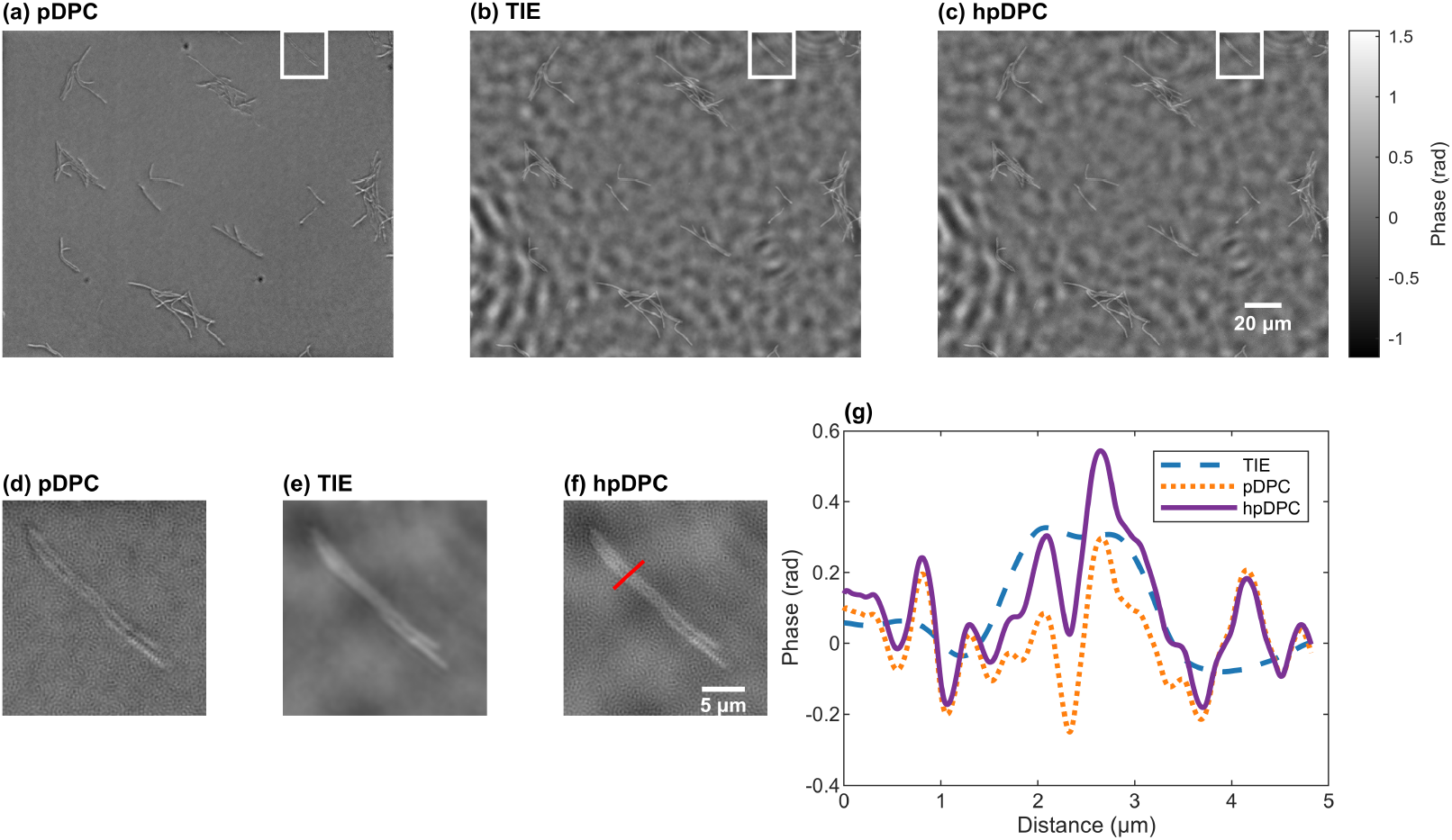
Comparison of phase reconstructions of *M. smegmatis* acquired with a 40 × 0.75 NA objective lens using pDPC, TIE, and hpDPC. (a–c) Full field-of-view phase reconstructions obtained using pDPC, TIE, and hpDPC, respectively. (d–f) Magnified views corresponding to the boxed regions in (a–c). The red line in (f) indicates the location of the extracted phase profile shown in (g). (g) Extracted phase profiles comparing TIE, pDPC, and hpDPC reconstructions. Scale bars: 20 µm in (c) and 5 µm in (f). Phase in radians.

Quantitative line profiles in Fig. 8(g), extracted along the transect indicated in Fig. 8(f), further illustrate these differences. The transect intersects two adjacent bacilli, giving an expected two-peak phase profile. TIE fails to resolve the two features, yielding a single broadened peak, whereas pDPC resolves the two peaks but with reduced phase amplitude. hpDPC preserves the two-peak structure while restoring phase contrast.

### 5.5. Learning-Based Cell State from Phase

Phase reconstructions provide an informative input representation for supervised learning tasks in label-free microscopy [6], including fluorescence inference and cellular-state classification. This is useful because cellular condition may be inferred without fluorescent dyes or chemical staining, enabling label-free measurements with fewer handling steps, reduced phototoxicity, and biological perturbation. For *M. smegmatis*, this is relevant to label-free assessment of membrane integrity and cellular state, for which fluorescence-based assays typically require exogenous staining. While fluorescence-based membrane-integrity assays are widely used for this purpose [31, 32], phase-based representations encode intrinsic morphological and biophysical properties of cells [1] and may therefore contain information complementary to fluorescence-based indicators. We evaluated whether the broader spatial-frequency response of hpDPC improves downstream discrimination of cellular state relative to pDPC and TIE alone.

Brightfield and phase morphology of heat-treated and control *M. smegmatis* samples can appear qualitatively similar at 60 ×, such that membrane compromise is not readily identifiable by visual inspection of the phase images alone. To evaluate whether more subtle phase-derived differences could nevertheless be exploited computationally, a supervised binary classification task was defined to distinguish image patches containing membrane-compromised cells from patches containing only non-fluorescent cells. Three phase reconstruction modalities were evaluated independently: hpDPC, pDPC, and TIE. Each classifier received only a single-channel phase image, without fluorescence information or engineered features.

As shown in Fig. 9(b) and (d), fluorescence imaging provides a direct indicator of membrane compromise following staining with SYTOX Orange. In contrast, the corresponding hpDPC phase maps in Fig. 9(a) and (c) do not show an explicit membrane-permeability marker. The classification experiment therefore tests whether subtle information contained in the reconstructed phase distributions can be extracted using a convolutional neural network (CNN).

**Fig. 9.**
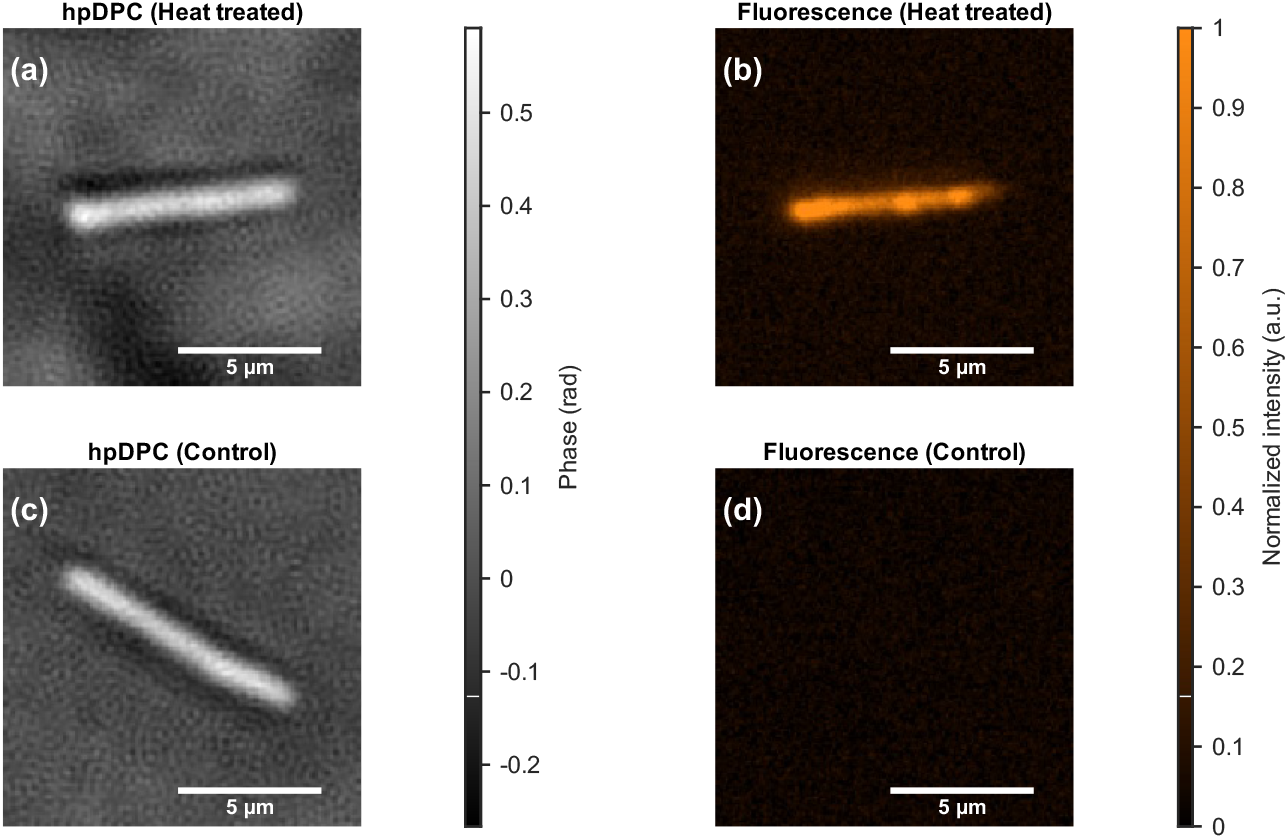
Matched hpDPC and fluorescence images of heat-treated and control *M. smegmatis* samples at 60 ×. (a,c) hpDPC phase maps and (b,d) corresponding fluorescence images for heat-treated (a,b) and control (c,d) conditions. Phase in radians. Fluorescence images are background-subtracted using the 5th percentile of control patches and normalized to the 99.5th percentile of the heat-treated signal for visualization. Scale bars: 5 µm.

#### 5.5.1. Dataset and Ground-Truth Labelling

The final analysis used 20 co-registered fields of view (FoVs), acquired using a 60 × 0.8 NA objective lens. Twelve FoVs were used for training, four for validation and model selection, and four for final evaluation. The same FoV assignments and fixed validation/evaluation coordinates were used for hpDPC, pDPC, and TIE to permit direct comparison between modalities. Each full-resolution phase image contained 2048 × 2448 pixels.

Binary cell masks were generated using the MATLAB Image Labeler app. SYTOX Orange fluorescence images were background-corrected by subtracting a Gaussian-blurred background (*σ* = 50 pixels), normalized, and denoised using non-local means filtering. Fluorescent regions were enhanced by morphological top-hat filtering (disk radius 5 pixels) and segmented by hysteresis thresholding, with high and low thresholds set to the 99.5th and 97th percentiles of the enhanced image, respectively. The resulting binary mask was morphologically cleaned and small connected components were removed to obtain the final SYTOX Orange segmentation mask. Phase and fluorescence images were co-registered using phase-correlation-based registration. Connected components were identified in the cell mask using 4-connected neighbourhood connectivity and designated membrane-compromised when any pixel overlapped the segmented SYTOX Orange mask.

Classification was performed at the patch level. Patch centres were sampled from foreground pixels in the cell mask, and 64 × 64-pixel phase patches were extracted around each coordinate. Candidate patches were accepted when at least 15% of their area was occupied by the cell mask and were labelled positive when any membrane-compromised connected component occurred within the patch; all others were labelled negative.

The training partition contained five heat-treated and seven control FoVs, while the validation and evaluation partitions each contained two heat-treated and two control FoVs. During each training epoch, 50 patches were requested from each of the 12 training FoVs, yielding 600 samples per epoch, with coordinates resampled at every epoch. Candidates with less than 15% cell-mask occupancy were rejected and resampled, so the final accepted contribution from individual FoVs could vary slightly. Validation and evaluation coordinates were generated once and held fixed across modalities and random seeds. Each set comprised 200 patches, with 50 requested from each of four FoVs. The resulting evaluation set comprised 100 positive and 100 negative patches; no class-balanced resampling was applied.

#### 5.5.2. Model Architecture and Training

A ResNet-18 [33] CNN was used independently for each phase reconstruction modality. The network was initialized from scratch rather than from pretrained weights. The original ResNet-18 stem was retained, except that the first 7 × 7 convolution was modified from three input channels to one input channel. Following the ResNet-18 global average-pooling layer, the original classification layer was replaced by a single linear unit mapping the 512-dimensional feature representation to one output logit. A sigmoid transformation was applied only when converting the output logit to a classification probability.

The phase patches were supplied as single-precision phase values without per-patch normalization or auxiliary channels. Separate hpDPC, pDPC, and TIE networks were trained for 200 epochs using Adam (10−3 learning rate; batch size 512) and weighted binary cross entropy with logits (positive-class weight 3.0). The class weight and fixed decision threshold of 0.3 were specified before final evaluation and applied identically across modalities; neither was optimized on the evaluation set.

To assess variability associated with stochastic initialization and patch sampling, each modality was trained independently using three random seeds (42, 43, and 44). For each seed, a fixed 200-patch validation set was evaluated after every training epoch. Model selection was performed exclusively using this validation set: the checkpoint achieving the highest validation F1-score at a fixed decision threshold of 0.3 was retained. No evaluation-set predictions were used for checkpoint selection in the final implementation. At inference, probabilities from the three seed-specific models were averaged to obtain the ensemble probability for each evaluation patch.

#### 5.5.3. Classification Performance

Classification performance was evaluated on the fixed set of 200 evaluation patches using receiver operating characteristic (ROC) and precision–recall analyses, summarized by the area under the ROC curve (AUC) and average precision (AP), respectively, together with F1-score, precision, and recall [34]. A fixed probability threshold of 0.3, identical to that used during validation-based checkpoint selection, was applied to all three modalities for threshold-dependent metrics. The threshold was not optimized separately for the evaluation data or for individual modalities. The resulting performance metrics are summarized in Table 1.

**Table 1.** Patch-level classification performance across phase reconstruction modalities on the fixed evaluation set.

| Modality | AUC | AP | F1 | Precision | Recall | TPR at 5% FPR |
| --- | --- | --- | --- | --- | --- | --- |
| hpDPC | 0.813 | 0.803 | 0.706 | 0.558 | 0.960 | 0.300 |
| pDPC | 0.672 | 0.619 | 0.667 | 0.500 | 1.000 | 0.050 |
| TIE | 0.444 | 0.449 | 0.648 | 0.492 | 0.950 | 0.000 |

Metrics were calculated from 200 fixed evaluation patches (100 positive and 100 negative) using the mean probability from three independently trained models. AUC and AP summarize ranking performance across decision thresholds. F1, precision, and recall were evaluated at a fixed probability threshold of 0.3; TPR at 5% FPR was obtained from the ROC curve. Patches were sampled from four FoVs. Abbreviations: AUC, area under the receiver operating characteristic curve; AP, average precision; TPR, true-positive rate; FPR, false-positive rate.

The EOC curves are shown in Fig. 10(a). The hpDPC ensemble achieved an AUC of 0.813, compared with 0.672 for pDPC and 0.444 for TIE. The separation was also evident in the low-FPR region: at an FPR not exceeding 5%, the corresponding true positive rate (TPR) values were 0.30 for hpDPC, 0.05 for pDPC, and 0 for TIE. The corresponding precision–recall (PR) curves are shown in Fig. 10(b). hpDPC achieved an AP of 0.803, compared with 0.619 for pDPC and 0.449 for TIE. At the fixed decision threshold of 0.3, hpDPC achieved a precision of 0.558, recall of 0.960, and F1-score of 0.706. pDPC yielded a precision of 0.500, recall of 1.000, and F1-score of 0.667, while TIE yielded a precision of 0.492, recall of 0.950, and F1-score of 0.648.

**Fig. 10.**
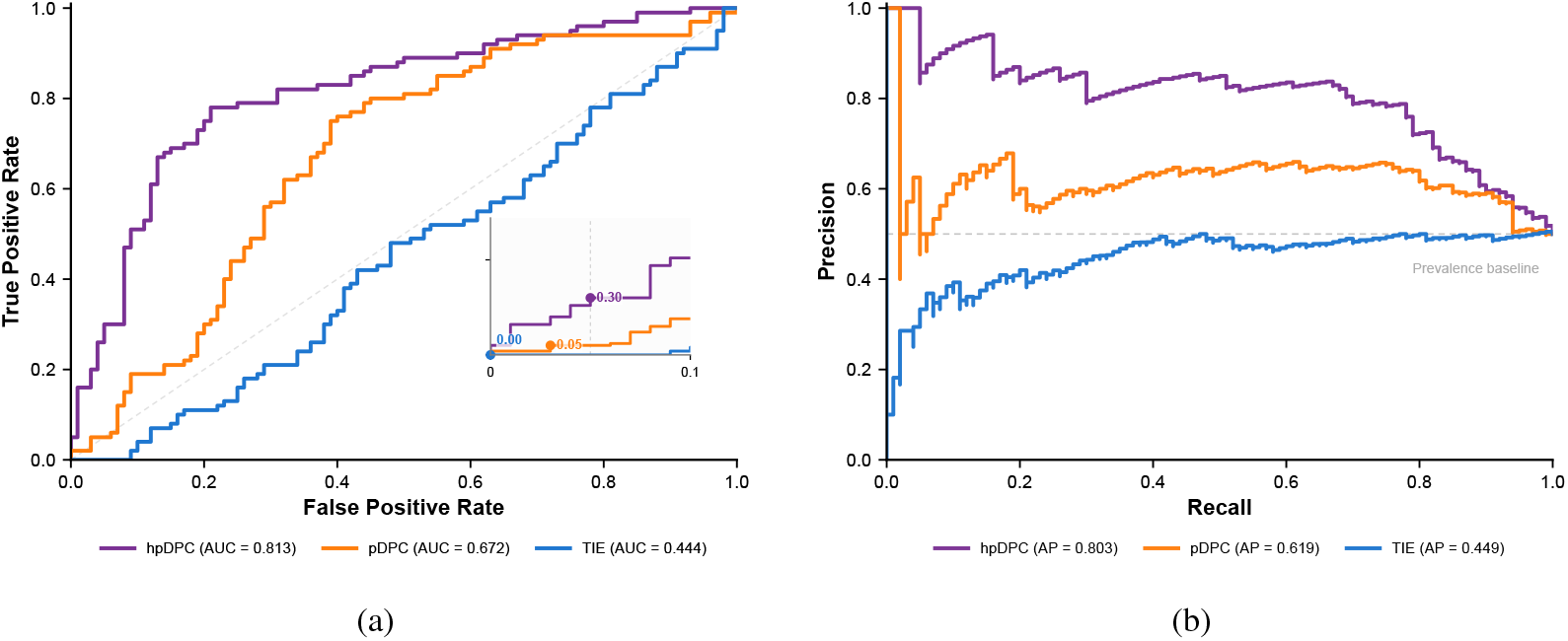
Patch-level classification performance for membrane-compromise inference using hpDPC, pDPC, and TIE phase reconstructions. (a) Receiver operating characteristic ROC) curves for the three-seed ensemble predictions. hpDPC achieved an area under the ROC curve (AUC) of 0.813, compared with 0.672 for pDPC and 0.444 for TIE. Inset: enlarged view of the ROC curves for false positive rate (FPR) < 0.1. (b) Precision–recall (PR) curves; corresponding average precision values were 0.803, 0.619, and 0.449 for hpDPC, pDPC, and TIE, respectively. The PR prevalence baseline reflects the 0.5 positive fraction of the fixed evaluation set.

## 6. Discussion

The spectral complementarity of hpDPC was evident in both simulations and experiments. pDPC preserved sharp boundaries and fine structure but underestimated slowly varying phase and phase amplitude, whereas TIE recovered smoother low-frequency phase at the expense of high-frequency detail. hpDPC combined these characteristics, improving phase recovery while retaining structural sharpness across the simulated microlens object, microspheres, A549 cells, and *M. smegmatis*. The approach also reduces the dependence of broadband phase recovery on strict matching between the pDPC illumination and objective NAs. Lower illumination NA can improve robustness to violations of the WOA, but reduces pDPC transfer support; in hpDPC, the resulting low-frequency loss is compensated by TIE, allowing pDPC to operate under more conservative illumination conditions.

The broader phase representation also improved downstream discrimination in the proof-of-concept classification experiment. hpDPC achieved an AUC of 0.813 and AP of 0.803, compared with AUC values of 0.672 and 0.444 for pDPC and TIE, respectively. At an FPR not exceeding 5%, hpDPC achieved a TPR of 0.30, compared with 0.05 for pDPC and 0 for TIE. These differences are consistent with attenuation of high spatial frequencies in TIE reducing sensitivity to fine structural variation, while reduced low-frequency sensitivity in pDPC may suppress broader morphological and optical-path-length information. Their combination therefore appears to provide a phase representation from which discriminative features are more readily learned. Given the limited number of independent FoVs, these results should be interpreted as proof-of-concept improved downstream discriminability rather than validation of a generalizable membrane-integrity classifier.

Polarization multiplexing also reduces acquisition burden by capturing the directional pDPC information in a single exposure. hpDPC therefore requires three intensity measurements: one pDPC frame and an in-focus–defocused TIE pair. This avoids motion between sequential asymmetric DPC measurements, although the TIE acquisition remains sequential and requires axial displacement. Simultaneous dual-plane detection or electronically controlled defocus could remove this remaining temporal separation. The polarization-sensitive detector further retains polarization-resolved intensity information. Although the present work considers only scalar phase, the acquisition architecture could potentially be extended to polarization-sensitive contrast in anisotropic specimens [35]. With appropriate calibration and polarization-aware forward modelling, established approaches [36] could be incorporated to estimate quantities such as retardance and optic-axis orientation. These capabilities were not evaluated here.

Several limitations remain. The reconstructed phase is semi-quantitative because both inversions rely on approximate forward models and regularization, while the single-sided finite-difference approximation used for TIE can introduce derivative error as the defocus distance increases. The Fourier fusion is also empirical: a fixed radial weighting is used rather than weights derived from calibrated transfer functions or spatial-frequency-dependent noise statistics, and no explicit inter-modality gain normalization is applied. Reconstruction accuracy may therefore depend on illumination calibration, polarization-channel balancing, and defocus accuracy. Future work should investigate transfer-function-and noise-informed fusion, quantitative calibration, simultaneous TIE acquisition, and validation across larger independent biological datasets.

## 7. Conclusion

We introduced hpDPC, a non-interferometric QPI framework that combines snapshot pDPC with TIE through co-designed illumination and frequency-selective Fourier-domain fusion. By combining the low-frequency sensitivity of TIE with the higher-frequency structural information recovered by pDPC, hpDPC provided broader effective spatial-frequency recovery than either modality alone while reducing the dependence of broadband phase recovery on strict illumination– objective NA matching. The framework requires only three intensity measurements and avoids sequential asymmetric DPC illumination through polarization multiplexing. Simulations and experiments with microspheres, mammalian cells, and *M. smegmatis* demonstrated improved recovery of both slowly varying phase and fine structural detail. Within a proof-of-concept phase-only classification task, hpDPC exhibited greater ROC and precision–recall discrimination than either constituent reconstruction (Fig. 10), indicating that the broader phase representation provided by hpDPC can also provide useful information for downstream computational analysis.

## Funding

This work was supported by the Wellcome Trust Grant 313779 and the Chan Zuckerberg Initiative Donor Advised Fund DAF2023-321240.

## Acknowledgment

The authors gratefully acknowledge Professor Paul M. W. French and Dr Sunil Kumar (LIGHT Community, Department of Physics, Imperial College London) for valuable discussions regarding pDPC microscopy and its implementation.

The authors thank Dr Dale Taylor, Holistic Drug Discovery and Development Centre, University of Cape Town, for providing the A549 cell cultures; Dr Alexa Rabeling, African Microscopy Initiative at the Institute of Infectious Disease and Molecular Medicine, University of Cape Town, for culturing the cells; and Isabel De Beer and the Molecular Mycobacteriology Research Unit at the Institute of Infectious Disease and Molecular Medicine, University of Cape Town, for providing the Mitomycin C-treated *M. smegmatis* samples.

## Disclosures

The authors declare no conflicts of interest.

## Data Availability

Data underlying the results presented in this paper, including simulation data, representative experimental data, processed phase reconstructions, and datasets used for quantitative analysis, are openly available in Zenodo at https://doi.org/10.5281/zenodo.22831229. Source code for phase fusion and simulation is available at https://github.com/frasermo/hpDPC-microscopy, with the version corresponding to this publication archived in Zenodo at https://doi.org/10.5281/zenodo.22962324.

## Supplemental document

See Supplement 1 for supporting content.

## Notes

### Competing Interest Statement

The authors have declared no competing interest.

